# GARFIELD - GWAS Analysis of Regulatory or Functional Information Enrichment with LD correction

**DOI:** 10.1101/085738

**Authors:** Valentina Iotchkova, Graham R.S. Ritchie, Matthias Geihs, Sandro Morganella, Josine L. Min, Klaudia Walter, Nicholas Timpson, UK10K Consortium, Ian Dunham, Ewan Birney, Nicole Soranzo

**Affiliations:** Human Genetics, Wellcome Trust Sanger Institute, Wellcome Trust Genome Campus, Hinxton, Cambridge, CB10 1HH, United Kingdom; European Molecular Biology Laboratory, European Bioinformatics Institute (EMBL–EBI), Wellcome Trust Genome Campus, Hinxton, Cambridge, CB10 1SD, United Kingdom; MRC Integrative Epidemiology Unit, University of Bristol, Oakfield House, Oakfield Grove, Clifton, Bristol, BS8 2BN, United Kingdom; Department of Haematology, University of Cambridge, Cambridge CB2 0AH, United Kingdom; The National Institute for Health Research Blood and Transplant Unit (NIHR BTRU) in Donor Health and Genomics at the University of Cambridge.

**Author notes:** Correspondence to: **Nicole Soranzo** Wellcome Trust Sanger Institute, Hinxton, CB10 1HH, UK, E-mail., **Ewan Birney**, The European Bioinformatics Institute (EMBL-EBI), Hinxton, CB10 1SD, UK.

## Abstract

Loci discovered by genome-wide association studies (GWAS) predominantly map outside protein-coding genes. The interpretation of functional consequences of non-coding variants can be greatly enhanced by catalogs of regulatory genomic regions in cell lines and primary tissues. However, robust and readily applicable methods are still lacking to systematically evaluate the contribution of these regions to genetic variation implicated in diseases or quantitative traits. Here we propose a novel approach that leverages GWAS findings with regulatory or functional annotations to classify features relevant to a phenotype of interest. Within our framework, we account for major sources of confounding that current methods do not offer. We further assess enrichment statistics for 27 GWAS traits within regulatory regions from the ENCODE and Roadmap projects. We characterise unique enrichment patterns for traits and annotations, driving novel biological insights. The method is implemented in standalone software and R package to facilitate its application by the research community.

## Introduction

Genome–wide association studies (GWAS) have discovered susceptibility variants for complex diseases and biomedical quantitative traits, with over 16 000 genotype–phenotype associations found to date ^1,2^, representing a large investment in resources, time and organisation to understanding human disease and other phenotypes. Despite the statistical soundness of the discovered associations, a large proportion (~90%) of implicated variants are classified as intronic or intergenic ^3^ and thus do not have a straightforward link to a cellular or molecular mechanism. This has prompted a number of efforts to annotate their putative functional consequences in cell specific contexts from experimentally derived regulatory genomic regions (e.g. regions marked by histone modifications, of open chromatin and transcription factor binding ^3–6^), principally as a means to inform and accelerate functional validation efforts.

The robust identification of which combinations of cells and marks are most informative for a given disease or quantitative trait of interest (henceforth referred generically to as 'phenotype') requires that one can confidently identify biologically meaningful correlations. Genomic marks may cover a large proportion of the genome, and thus many disease–associated variants will be found within these marks by chance. In addition, the heterogeneous distribution of genetic variants and functional regions along the human genome, and thus non–random association with genomic features ^7,8^, can create spurious correlations that again confound correct interpretation.

Functional enrichment methods exploit experimentally derived regulatory genomic regions to assess the relative contribution of variation in each cell type and regulatory annotation to a given phenotype of interest. In their simpler implementation, they estimate enrichment of association p–values based on comparisons of the full set of genetic variants analysed in the GWAS study ^9–11^, or on subsets of highly associated variants, for instance variants achieving genome–wide significance ^12–14^. These approaches have identified many biologically plausible patterns of correlation (for instance in open chromatin marks for lipid traits in liver cell types and Crohn’s disease in immune cells) and are broadly used for ranking the relative contribution of features. However, there is currently little confidence in interpreting unexpected enrichment, because of concerns with the statistical methodology, for three main reasons. Firstly, overly simplistic genetic models that do not account for known confounders such as local linkage disequilibrium (LD), local gene density and minor allele frequency, can lead to spurious enrichment patterns ^12^. Second, reliance on restrictive parametric statistics (rather than permutations) makes these approaches less robust to the well–established underlying heterogeneity of genomic feature distribution. Finally, tests based on subsets of variants (for instance those reaching genome–wide significance) typically probe a limited number of genomic features, while it has been shown that evidence of enrichment occurs well below genome–wide significance ^9,10^. Methodological improvements are thus needed to improve the accuracy of inference, and to realise the full potential of those costly experiments in focused experimentation.

Here we present a novel nonparametric approach that leverages GWAS findings with regulatory or functional annotations to find features relevant to a phenotype of interest. To our knowledge, this is the only method that accounts for LD, minor allele frequency, matched genotyping variants and local gene density with the application of permutations to derive statistical significance. We name our method **GARFIELD**, which stands for **G**WAS **A**nalysis of **R**egulatory or **F**unctional **I**nformation **E**nrichment with **LD** correction. We used GARFIELD to analyse the enrichment patterns of publicly available GWAS summary statistics using regulatory maps from the ENCODE ^3^ and Roadmap Epigenomics ^5^ projects. Finally, we developed new software to facilitate the application of our approach by the research community, and tools for effective visualisation of enrichment results that scale to thousands of potential functional elements. In our own use we have discovered expected and novel enrichments that illustrate the molecular and cellular basis of well studied traits, and we expect this method to help drive novel biological insights and enhance efforts to robustly prioritise variants for follow–up studies across existing and future association studies.

## Results

### Overview of the method

The analysis workflow implemented in GARFIELD is summarised in Figure 1 and Online Methods. The method requires four inputs: (i) a set of genome–wide summary statistics, corresponding to single–variant p–values for the association of genetic variants with a given disease or trait of interest; (ii) genome–wide genomic coordinates for a regulatory feature of interest; (iii) a list of LD tags for each variant (measured by the r^2^ statistic with values of 0.1 and 0.8 within 1MB windows) from a reference population of interest (e.g. Caucasian) and (iv) minor allele frequency (MAF) and distance to the nearest transcription start site (TSS) measurements. Given these inputs, the method performs the following steps: (i) it greedily reduces the genome–wide genetic variants to an independent set using LD and distance information (‘LD pruning step’) by sequentially removing variants with r^2^>0.1 and within 1Mb window from the most significantly trait–associated variant; (ii) it annotates each variant with a regulatory feature if either the variant, or a correlated variant (r^2^>0.8), overlaps the feature (‘LD tagging annotation step’); (iii) and finally it calculates fold enrichment (FE) at different GWAS p–value thresholds (denoted as ‘T’) and tests their significance by comparing the observed enrichment to the one based on a large number of permutations for each annotation while performing ‘feature matching’ (Online Methods) on variants by MAF, distance to the nearest TSS and number of LD proxies (r^2^>0.8). To correct for multiple testing on the number of different annotations, it further estimates the effective number of independent annotations by using the eigenvalues of the correlation matrix of the binary annotation overlap matrix from Figure 1 (adapted from Galwey et.al. 15) (Online Methods; Supplementary Figure S1) and then applies a Bonferroni correction at the 95% significance level. This takes into account the tissue selective components of regulatory data, namely that closely related cell types and tissues are more similar to each other than different ones.

**Figure 1.**
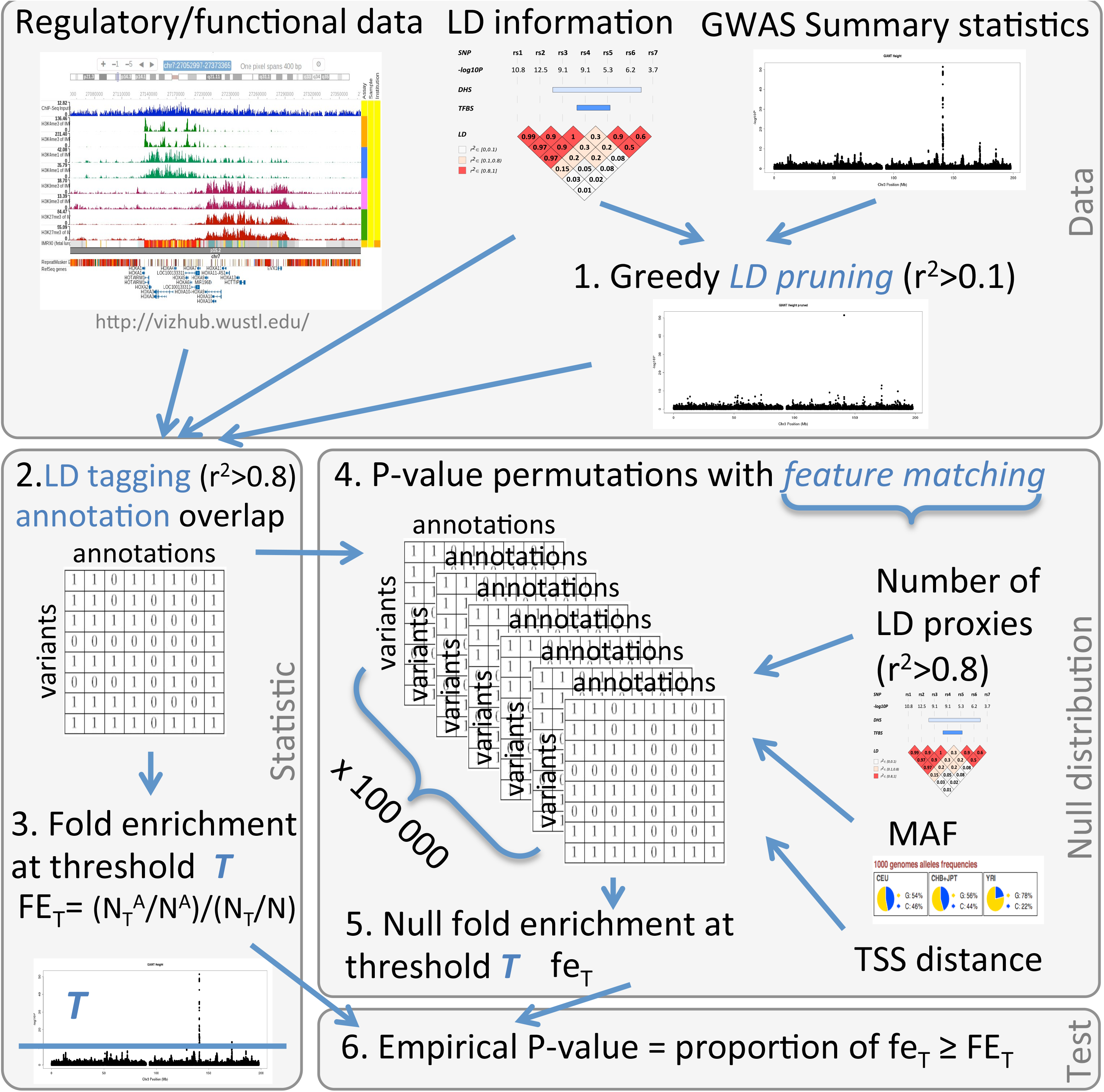
GARFIELD method flow. The ‘Data’ panel shows the input annotation, p–value and LD data and the first step of LD pruning. The ‘statistic’ panel presents (i) the binary annotation overlap matrix for all pruned variants and all annotations (with overlap denoting physical overlap or LD r^2^>0.8 based one); (ii) The fold enrichment statistic at a GWAS significance P–value threshold T, where N denotes the total number of independent variants, NT– the subset of them with P–value less than T, NA– the number of variants in the annotation of interest, and NTA– the number of them with P–value less than T. The ‘Null distribution’ panel describes the permutation procedure for producing a null distribution for our test statistic. Namely, it involves a large number (e.g. 100 000) of permutations of the p–values of the variants in our independence set, by feature matching to MAF, TSS distance and number of LD proxies. Finally, panel ‘Test’ shows the empirical enrichment p–value calculation.

Our approach can be viewed as similar to Maurano et.al. ^9^ (see also Supplementary Table S1) with two critical improvements. First, we account for the effect of local correlation between variants by restricting FE calculations to sets of independent variants (LD pruning step). Second, we employ an adaptive permutation procedure that creates and utilizes null variant sets that account for systematic differences in MAF, gene distance and number of proxies in the test variant set. Figure 2A illustrates the FE and significance results under GARFIELD and two simpler approaches (NM = naive model with no LD or genomic feature correction (corresponding to the Maurano model); LDM = LD–pruned model with no LD tagging annotation or feature correction) for the Crohn’s disease phenotype (at T<10–^8^) in DNaseI hypersensitive sites (ENCODE, Roadmap Epigenomics) for annotations that showed significant enrichment in at least two of the methods. Results revealed a significant decrease in FE estimates from NM to LDM and GARFIELD models, a result of potentially multiple genetic variants tagging a single underlying trait association.

**Figure 2.**
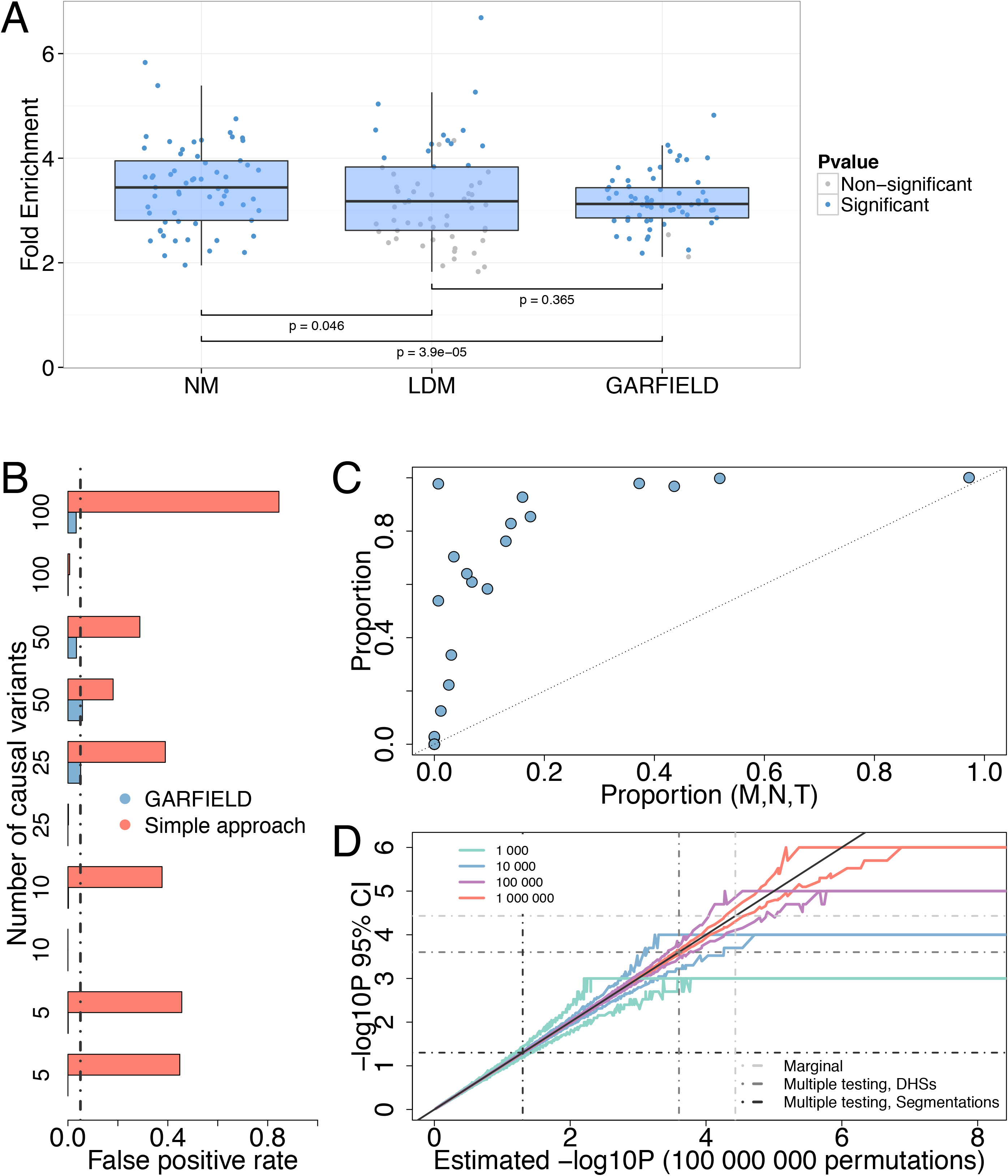
Method assessment. (A) FE and significance of enrichment for an example trait, Crohn’s Disease (CD) for DNaseI hypersensitive sites in 424 cell types, from ENCODE and Roadmap Epigenomics, for annotations with significant enrichment in at least two methods out of NM, LDM and GARFIELD. Between method p–values were obtained using a paired Wilcoxon two–sample test. (B) Estimated false positive rate (FPR) for 10 simulated datasets containing different numbers of causal variants (y–axis) and 1000 simulated annotations. Black vertical line denotes the 5% FPR threshold. (C) Comparison between the proportion of significant annotations found from models accounting for MAF (M), number of proxies (N) and distance to nearest TSS (T) respectively, to a model not accounting for any feature, for each of 27 publicly available GWA studies and 424 DNaseI hypersensitive site annotations. (D) Estimates of 95% confidence intervals for–log10 enrichment P– value for the Crohn’s Disease trait and DNaseI hypersensitive site data based on 1000 runs of different numbers of permutations (from 104 to 107). Horizontal coloured line segments denote the minimum non–zero P–value that can be obtained after n permutations and represent p less than or equal to this value (p=0 is denoted as equal to 1/n for the analyses). Dotted lines represent 5% significance thresholds for marginal tests and tests after multiple testing correction for the DHS and segmentation data to be used in subsequent analyses.

To assess these models further, we simulated genome–wide association summary statistics for 10 quantitative phenotypes with additive association to 5–100 randomly selected genetic variants (0.3<beta<1, MAF >5%) (Methods). For each of them, we estimated FE metrics under the NM and GARFIELD models against 1000 peak region annotations, simulated to match observed peak lengths and between peak distances for DNaseI hypersensitive sites in HepG2 cells (ENCODE). False positive rate (FPR) estimates were then obtained by calculating the observed proportion of significantly enriched annotations per phenotype. As a result, we found a significant increase in FPR when LD is not explicitly modelled (from an average of 0.02 for GARFIELD to 0.3 for NM at the 5% significance level, Wilcoxon two–sample paired test p–value= 9x10^-3^) (Figure 2B).

Additionally, to assess the value of feature matching in significance testing, we employed GARFIELD with and without MAF, TSS distance and number of LD proxy correction to 424 open chromatin annotations in 27 phenotypes. As expected, we found that feature matching further controls for biases in enrichment analysis by significantly reducing the number of observed significant enrichments (Wilcoxon signed rank test proportion median–0.50, p–value = 1x10^-4^) (Figure 2C). We further explored the relative contribution of each feature in turn by comparing GARFIELD’s results when correcting for it against those with no feature correction and found median proportion reduction estimates of significant enrichments of–0.37 (p–value =1x10^-4^),–0.11 (p– value =7x10^-4^) and 0.02 (p–value = 2.3x10^-3^) for the number of LD proxies, TSS distance and MAF, respectively (Supplementary Figure S7). These tests highlight the number of LD proxies as the most important single confounder, however not sufficient to correct for individually when compared to using all three features together.

Finally, we estimated the uncertainty in the observed enrichment P–values by running GARFIELD 1000 times with each of 10^3^, 10^4^, 10^5^ and 10^6^ permutations for the Crohn’s disease phenotype in 424 open chromatin annotations. As expected the larger the number of permutations, the tighter the confidence intervals around estimates, which however comes at a higher computational cost (Figure 2D). In practice we use 10^5^ permutations for the open chromatin data and 10^6^ for the genomic segmentations which results in accurate estimation of enrichment P–values up to ~1x10^-4^ after multiple testing correction. Additionally, we implement an adaptive permutation procedure which terminates the iterations if significant enrichment of a given feature can no longer be achieved (after minimum of 100 iterations). This provides a substantial reduction in method runtime (from an average of 20.6 min to 1.8 min per traits for the open chromatin data at the 10^-8^ GWAS threshold) (Supplementary Figures S6).

### Enrichment in open chromatin regions from 424 cell types

To assess the relative enrichment of phenotype–genotype associations in different cell types, we first applied GARFIELD to a generic open chromatin mark, DNase I hypersensitive sites, commonly used as a marker of regulatory DNA, in 424 cell lines and primary cell types from ENCODE 3 and Roadmap Epigenomics ^5^ (Supplementary Table S2). We considered 3 disease and 24 quantitative traits with publicly available GWAS summary statistics. For each trait and annotation pair we derived FE statistics at eight GWAS P–value thresholds (T<10^-1^ to T<10^-8^). At the most stringent cut–off (T<10^-8^), there were a median of 19 independent SNPs tested genome–wide per trait (range 0^-331^, Table 1 and Supplementary Table S3), while selecting SNPs using a more permissive threshold (T<10^-5^) increased the number of SNPs tested to a median of 77 variants per trait (range 11^-707^).

**Table 1.**
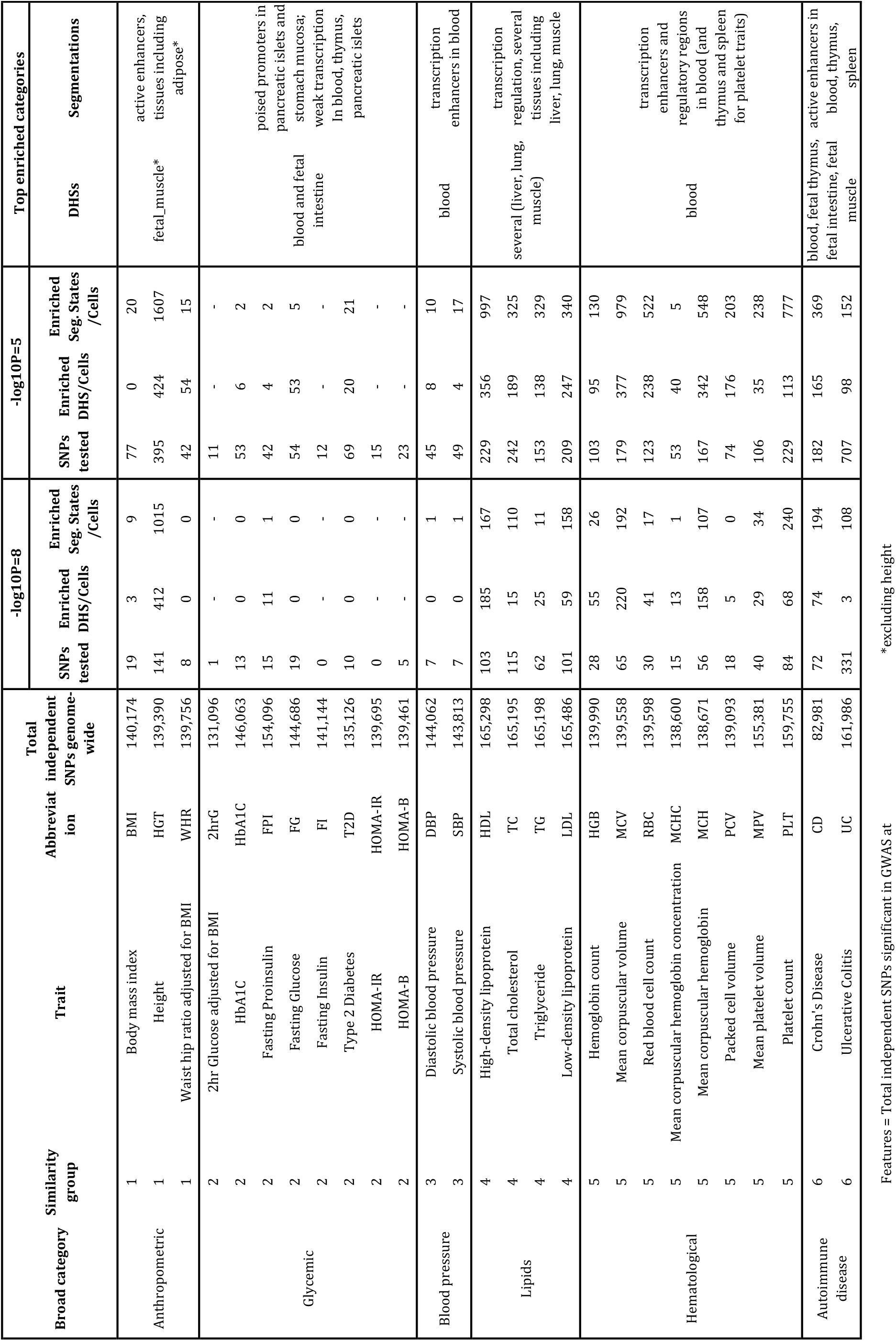
Summary of enrichment analyses in DNaseI hypersensitive sites and genomic segmentations per phenotype.

We further tested the significance of observed FE statistics at the four most stringent thresholds (T<10^-5^ to T<10^-8^). We found statistically significant enrichments (defined by an empirical p<2.6x10^-4^, see Online methods) for the majority of traits considered, highlighting clear differences in enrichment patterns between traits (Supplementary Table S4). As also clearly visible from enrichment wheel plots, some traits displayed relatively ubiquitous enrichment (e.g. height in Figure 3b), as compared to traits with relatively narrow enrichment (e.g. Crohn’s disease, Figure 3a, see also Supplementary Figures S3–S5). Blood cells were overall the most enriched tissue type in all haematological traits and autoimmune diseases, but provided little to no enrichment for glycemic, blood pressure and anthropometric traits, with the exception of height which was enriched in nearly all tissues. As predicted, incorporating sub–threshold associations (T less than 5x10^-8^) increased the number of variants added to the analysis, which in turn greatly increased the resolution of enrichment patterns across different traits (Table 1). For instance, at T<10^-8^ there were no annotations enriched for WHR (waist–to–hip ratio), while at the more permissive threshold T<10^-5^ there were 54 significant enrichments, mostly corresponding to muscle or fetal muscle tissue. For Ulcerative Colitis (UC), there were three annotations enriched at 10^-8^, but a much larger number (98) at 10^-5^, including several blood cell types and interestingly also foetal intestinal tissue.

**Figure 3.**
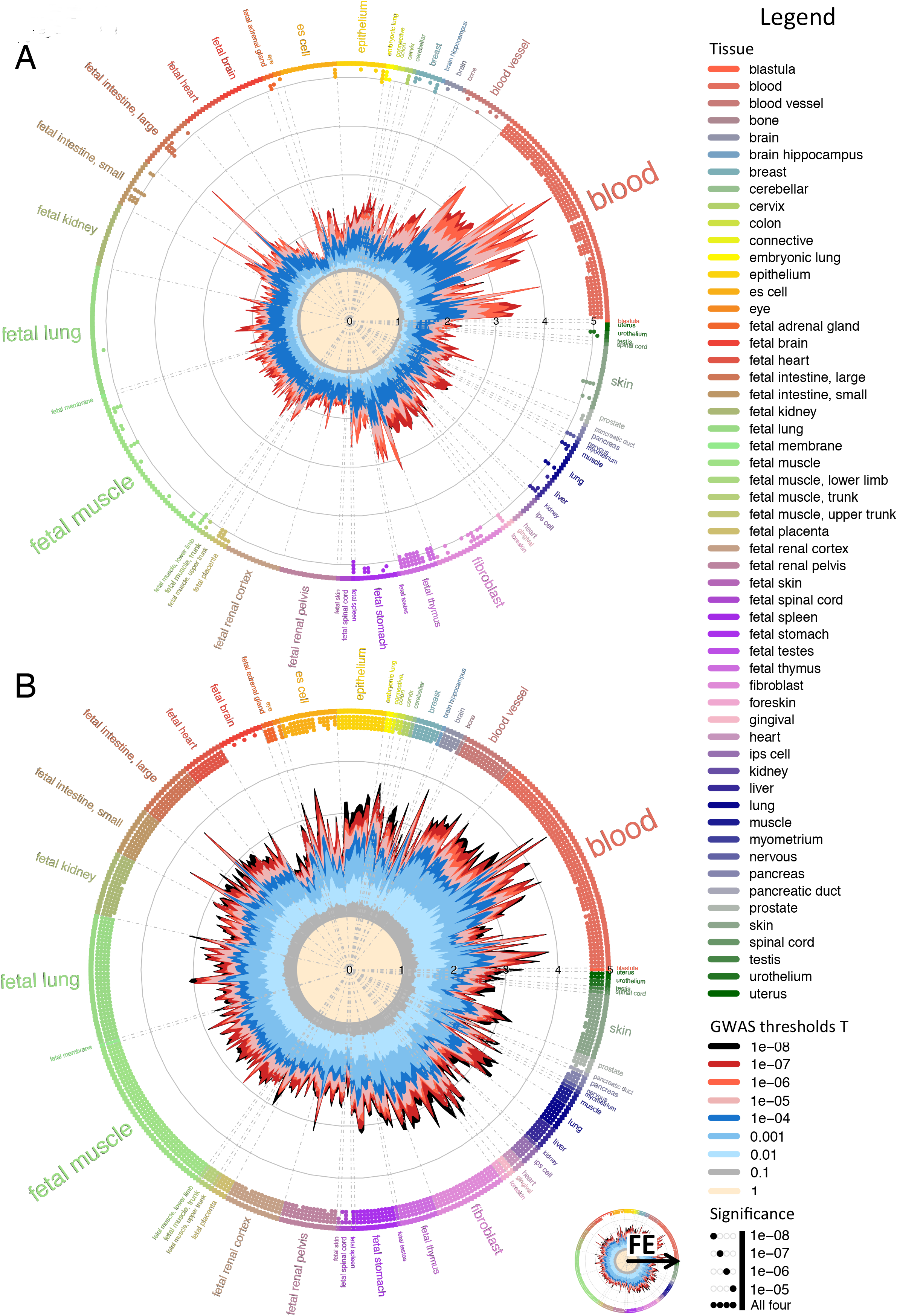
Enrichment of genome–wide association analysis p–values in DNaseI hypersensitive sites (hotspots). (A) Crohn's disease (CD). (B) Height. Radial lines show FE values at eight GWAS P–value thresholds (T) for all ENCODE and Roadmap Epigenomics DHS cell lines, sorted by tissue on the outer circle. Dots in the inner ring of the outer circle denote significant enrichment (if present) at T<10^-5^ (outermost) to T<10^-8^ (innermost) and are coloured with respect to the tissue of the cell type they test. CD shows to be predominantly enriched in blood, fetal thymus and fetal intestine tissues whereas HGT exhibits an overall well spread enrichment.

These enrichments reflect current understanding of key cellular types for disease, augmented with novel observations. In the former category were enrichments of lipid traits in blood, liver, fetal intestine and fetal thymus cell types; of haematological traits in blood and blood vessel tissues, and of autoimmune diseases (UC and CD) in blood and fetal intestine 9,11,16. An interesting example of an observation not previously described is the enrichment of Caco–2 (a well established gut epithelia cellular model) elements for LDL (Low Density Lipoprotein) and TC (Total Cholesterol), surpassing that for the expected key enrichment in HepG2 (a well established hepatocyte cellular model) (FE=3.4 in Caco–2 and FE=2.2 in HepG2 for LDL for T<10^-8^). Underlying, 52% of Caco–2 DNaseI peaks are shared with HepG2 (>=1bp overlap) and 36% of UK10K^17^ sequence variants overlapping Caco–2 also overlap HepG2 (Supplementary Figure S9) (average of 12 LD tags per variant). Furthermore 68% of Caco–2 annotated independent LDL associated variants were also shared with HepG2 (average of 11 LD tags per variant) and this proportion increased further when looking at the T<10^-8^ threshold to 74%, representing a much larger extent of sharing than expected from the DNaseI or genotype data alone. Unsurprisingly, we found the genes close to shared Caco–2/HepG2 LDL associated variants to be associated with lipid functions, as expected from the known cholesterol pathways in liver (GREAT GO enrichment analysis 18, Supplementary Table S5). Interestingly however, when performing GO enrichment analysis on the subset of LDL associations overlapping Caco–2 DNaseI peaks, but not HepG2 ones, we found enrichment in T cell receptor function (Supplementary Table S5), which was not observed for shared or HepG2 specific variants and those not overlapping either HepG2 or Caco–2 DNaseI peaks. Together with the overall stronger enrichment of LDL variants in Caco–2 than HepG2, this suggests that there might be a different aspect of cholesterol management mediated by gut epithelia, potentially with the involvement of the immune system.

### Enrichment in genomic segmentations of 127 epigenomes

We additionally sought to compare the relative enrichment of different types of functional genomic marks, using data on genomic segmentations for 127 cell types (Supplementary Table S6), where twelve (imputed) marks were used to obtain a 25–state model (Supplementary Table S7) using ChromHMM. For each segmentation state and cell type we analysed the same 27 phenotypes investigated before at 4 GWAS p–value thresholds (T<10^-5^ to T<10^-8^). Overall, when considering only significantly enriched trait–annotation pairs (defined by an empirical p<3.7x10^-5^), we found higher levels of FE for promoters (median 8.7, range [5.9^-13^.8] for T<10^-5^) and enhancers (median 7.4, range [4.3^-12^.5]) as opposed to transcribed regions (median 4.0, range [2.8^-7^.1]) (Figure 4C) (similar patterns were obtained for T<10^-8^, Supplementary Figure S13).

**Figure 4.**
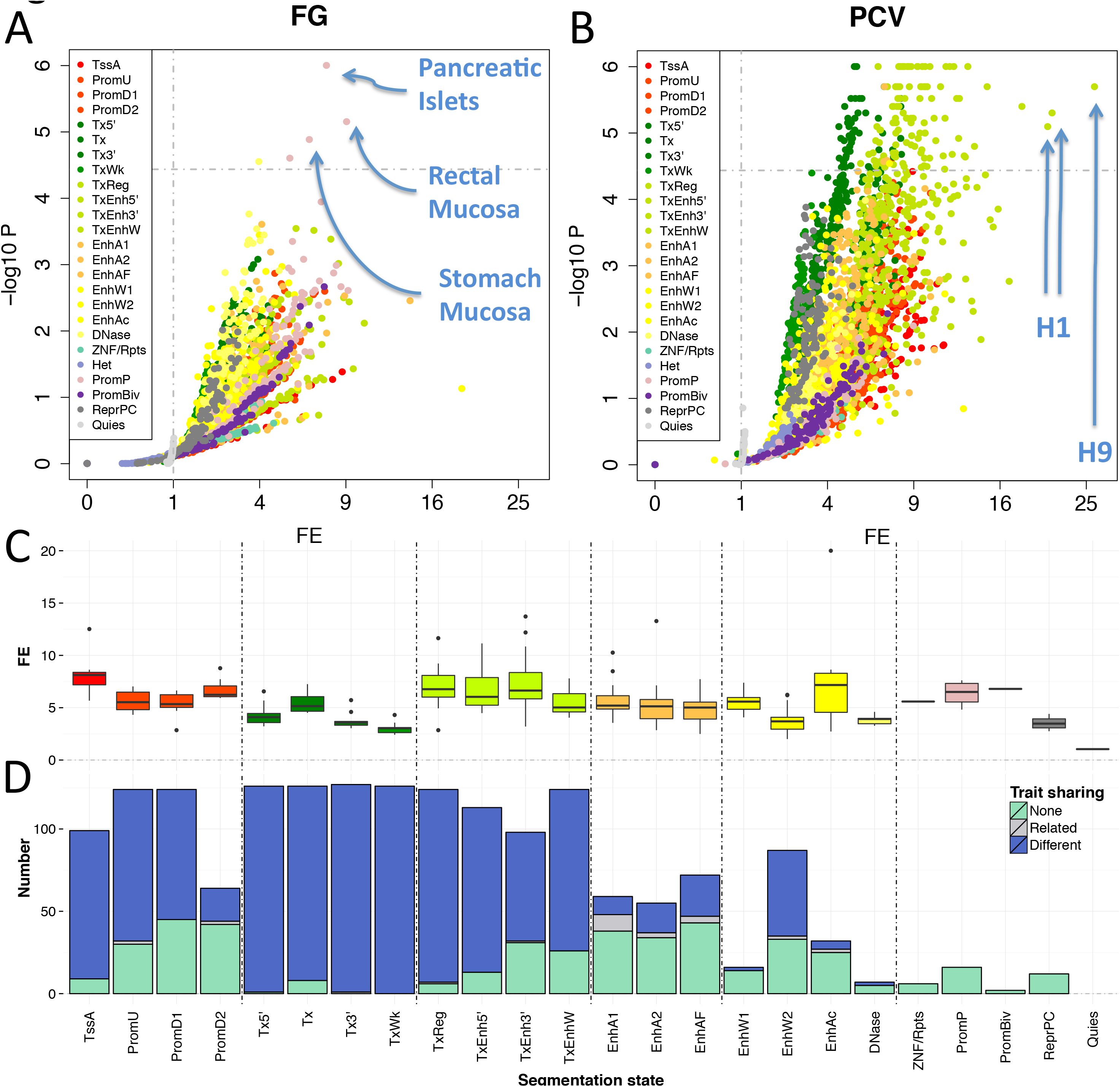
Fold enrichment levels and extent of sharing between traits for 25–state chromatin segmentations of the NIH Roadmap and ENCODE projects at the T<10^-5^ GWAS significance threshold. (A–B) Example feature volcano plots for Fasting Glucose (FG) and Packed cell volume traits (PCV), respectively. (C) Distribution of significant FE values across the 27 traits considered, split by segmentation state and coloured to highlight predicted functional elements (e.g. bright red for active TSS, dark green for transcription, see Supplementary Table S7). (D) Sharing of significantly enriched annotations with FE>2 across different phenotypes. The barplot displays the number of cell types where an annotation is uniquely enriched in a trait (light green), shared between closely related phenotypic traits (e.g. LDL cholesterol and TC, see Supplementary Table S3) (grey) or shared among non–correlated traits (e.g. TC and height) (blue).

The majority of these were found to come from transcription (30-32%) and transcription enhancer states (21– 25%) followed by weak transcription (12%) and downstream or upstream promoters regions (13-15%). This could be a result of larger power to detect significant enrichment for transcription states due to their larger region size. In terms of cell type specificity, similarly to the open chromatin data, we see the trait height as the most ubiquitously enriched phenotype. In general, we find the largest FEs for anthropometric traits in active enhancers in adipose tissue; glycemic traits in poised promoters in pancreatic islets and stomach mucosa and weak transcription in blood, thymus and pancreatic islets; lipid traits in transcription regulation in tissues including liver, lung and muscle; autoimmune diseases and platelet traits in active enhancers in tissue including blood thymus and spleen. As expected, incorporating sub–threshold associations again greatly increases the resolution of enrichment patterns across different traits (Table 1). For example, we find no significant enrichment at T<10^-8^ for Type 2 diabetes (T2D), fasting glucose (FG), fasting proinsulin (FPI) and packed cell volume (PCV), whereas at T<10^-5^ T2D is enriched in weak transcription and weak enhancer state in primary T helper cells, primary B cells, pancreatic islets and thymus, FG in poised promoters in pancreatic islets, stomach and rectal mucosa, PCV predominantly in transcription regulation in blood and es cells.

We further assessed the extent to which traits identifying closely related functions (e.g. anthropometric, hematological, glycemic, lipid, blood pressure and autoimmune disease; for short denoted as *related* or *similar* traits; Table 1), shared significantly enriched annotations as opposed to non–related traits (Supplementary Figures S9–S11 and Supplementary Table S8). To do this we counted the number of cell types per segmentation state that were found to be significantly enriched (and have FE greater than 2) for (i) only a single trait, (ii) at least two closely related traits (and no non–related ones), (iii) at least two non–related traits (Figure 4D). As a result we found higher cell type specificity for promoter and enhancer states (average percentage of enriched cell types unique to a single trait of 34% for promoters and 65% for enhancers), and larger sharing for transcribed regions (average of 9% being trait specific). Similarly, the percentage of cell types shared between similar traits from those shared between any type of traits was largest for enhancers, followed by promoters and smallest for transcribed states (average of 18% for enhancers, 3% for promoters, 0.4% for transcribed regions). This higher sharing for transcribed regions could reflect underlying biological processes, but may also be due to the larger expected power for enrichment detection in broader annotations. Finally, we observed a lot less sharing between related traits than that including non–related traits with the exception of active enhancers (EnhA1 with 47% of sharing occurring between similar traits), which we suspect to be due to the smaller sizes of classes of related traits when compared to the total number of trait classes.

### Software implementation

Functional enrichments analyses have been part of many GWA studies, which have been often performed with customized, in–house pipelines. In order to facilitate the application of our approach by the research community, we implemented GARFIELD as a standalone tool in C++ (Online Methods). The software allows for enrichment analysis of any user–provided trait with the following required input information: variant GWAS p– values and genomic coordinated in build37. We further provide over 1000 GENCODE ^19^, ENCODE ^3^ and Roadmap Epigenomics ^5^ pre–compiled annotations, UK10K sequence LD data and MAF and TSS distance informations for a ready to use package. Furthermore, custom user annotation data can also easily be accommodated when provided in a simple bed format. The tool consists of two main parts: (i) pruning and annotation of the GWAS study of interest and (ii) calculating fold enrichment and testing its significance. Memory and CPU usage information for both is given in Supplementary Table S9 and Supplementary Figure S6. As well as this standalone software we have also developed a Bioconductor package for the R statistical framework to further ease usability.

## Discussion

Large–scale efforts, such as those for ENCODE and Roadmap Epigenomics projects, have been devoted to systematically mapping molecular traits associated with regulatory genomic regions. They have greatly enhanced the annotation of putative functional consequences of non–coding variants in cell specific contexts, and have further shown to provide links to disease association. However, current methods that aim to evaluate the contribution of such regions to genetic variation in disease cannot always do so robustly or are not readily applicable for systematic analysis and comparison of broad sets of features. In particular, it has been shown that LD, gene density and MAF can confound enrichment analysis results ^12^. Here we further estimated the relative effect of each of those features and identified LD as the largest confounder. Additionally, by design, different genotyping platforms can create different biases (e.g. number of variants, allele frequency distribution, genomic location distribution). GARFIELD accounts for all those features and to the best of our knowledge there is no other method that can do so without making extremely restrictive assumptions (e.g. Pickrell et.al. ^11^ assume at most one causal variant at a given genomic region). Furthermore, the majority of available approaches typically use variants that reach genome–wide significance from association analysis (T<5x10^-8^) although there has been evidence of enrichment occurring well below that level ^9,10^. To capture these effects, GARFIELD allows for parallel enrichment analyses at multiple p–value sub– thresholds, which improves power to define statistically significant enrichment patterns by increasing the number of variants tested, thus enabling its application to traits with underpowered GWAS studies. Finally, we provide a flexible software platform with effective visualisation to enable researchers to carry out simultaneous enrichment analysis for thousands of annotations at multiple association thresholds.

In our own application of GARFIELD on existing GWAS and functional datasets we identified a broad set of largely expected or previously identified enrichments, for example lipids traits in open chromatin in liver, haematological traits in blood and anthropometric traits in active enhancers in adipose tissue. We further discovered a much larger enrichment of LDL cholesterol in open chromatin regions in colon cells (Caco–2) than in liver (HepG2), an interesting example, potentially related to lipid uptake by the gut epithelia. A number of GWAS hits do not show significant enrichments even with established cell types when using higher thresholds, but GARFIELD’s progressive, stratified approach uncovers these more nuanced enrichments, shown in the case of pancreatic islets with T2D. By analysing large–scale genome segmentation data, we assessed the relative contribution of each segmentation state to the phenotypic traits. We discovered a larger number of enrichments coming from transcription states as opposed to promoter and enhancer states together with a larger number of shared cell types between traits. These findings may be biologically relevant but they could also be a result of statistically larger power for enrichment detection for broader region annotations. On the other hand, we observed (~2 fold) larger FE values for significant enrichments in promoter and enhancer regions when compared to those for transcribed region, highlighting them as much more relevant for trait associated variants.

Robust, usable and modular methods are critical in the modern large–scale analysis arena, where we expect many discoveries to come from principled combinations of heterogeneous datasets. We have already deployed the GARFIELD method in a number of association study settings both in house and more broadly in the community. Our aim in developing GARFIELD has been to provide the most robust statistical framework for analysing functional enrichments coupled with practical ease of use and visualisation, and we hope the community will continue to exploit this tool to provide more insights into disease mechanisms.

## Acknowledgements and grant support

This study makes use of data generated by the UK10K Consortium.. A full list of the investigators who contributed to the generation of the data is available from www.UK10K.org. Funding for UK10K was provided by the Wellcome Trust under award WT091310. NIcole Soranzo's research is supported by the Wellcome Trust (Grant Codes WT098051 and WT091310). NJT works within the MRC Integrative Epidemiology Unit (Medical Research Council MC_UU_12013/3); EMBL–WTSI (G.R.S. Ritchie, V. Iotchkova).

## Author Contributions

Contributed data or materials: G.R.S.R., J.L.M., K.W., N.J.T., I.D., N.S.; Developed the method: E.B., G.R.S.R., I.D., J.L.M., N.S., V.I.; Analysed the data: V.I.; Provided critical interpretation of results: E.B., I.D., N.J.T., N.S., V.I.; Designed tools: M.G., S.M.; Wrote the manuscript: E.B., N.S., V.I.; Evaluated the manuscript: E.B., G.R.S.R., I.D., J.L.M., K.W., M.G., N.J.T., N.S., S.M., V.I.; Designed and managed the project: E.B., N.S.

## Online Methods

### Association Summary Statistics Data

GWAS summary statistics from the analysis of 27 disease and quantitative phenotypes were obtained from a number of sources. From GIANT (http://www.broadinstitute.org/collaboration/giant/index.php/GIANT_consortium) we downloaded large studies on BMI^20^, Height^21^ and Waist hip ratio adjusted for BMI^22^. From MAGIC (http://www.magicinvestigators.org/downloads) we downloaded data on BMI adjusted 2hr glucose^23^, HOMA B, HOMA IR, Fasting glucose, Fasting insulin^24^, Fasting proinsulin^25^ and HbA1C^26^. Global lipid GWAS summary statistics for LDL, HDL, TC and TG^27^ were obtained from (http://www.sph.umich.edu/csg/abecasis/public/lipids2010/). IIBDGC data on Crohn’s disease^28^ and Ulcerative colitis^29^ was obtained from (http://www.ibdgenetics.org/downloads.html). ICBP data on SBP and DBP^30^ was downloaded from (http://www.georgehretlab.org/icbp_088023401234-9812599.html). Type 2 diabetes^31^ GWAS summary statistics were downloaded from DIAGRAM (http://diagram–consortium.org/downloads.html). Blood trait data on HGB, MCH, MCV, MCHC, RBC and PCV was additionally obtained from the authors of Soranzo et. al. ^32^ and MPV and PLT data from the authors of Gieger et. al.^33^(Supplementary Table S3).

### DHS data

DNaseI hypersensitive sites (hotspots) were obtained from ENCODE (http://genome.ucsc.edu/cgi–bin/hgTrackUi?db=hg19&g=hub_4607_uniformDnase&hubUrl= http://ftp.ebi.ac.uk/pub/databases/ensembl/encode/integration_data_jan2011/hub.txt) and the NIH Roadmap Epigenomics Mapping (http://www.genboree.org/EdaccData/Current–Release/experiment–sample/Chromatin_Accessibility/) on all available cell types. DHS data was processed following DHSs data processing protocol described in an ENCODE study^4^. Further information on the data can be found in Supplementary Table S2.

### Epigenome segmentation data

Data from a chromatin state model with 25 states based on imputed data for 12 marks (H3K4me1, H3K4me2, H3K4me3, H3K9ac, H3K27ac, H4K20me1, H3K79me2, H3K36me3, H3K9me3, H3K27me3, H2A.Z, and DNase) across 111 Roadmap Epigenomics^14^ and 16 ENCODE reference epigenomes was downloaded from http://egg2.wustl.edu/roadmap/web_portal/. State and cell line information can be found in Supplementary Tables S7 and S6.

### LD data

LD information (proxies) was calculated using PLINK^34^ (v1.7) and the––tag–r2 0.1––tag–kb 500 (and––tag–r2 0.8–– tag–kb 500) flags in order to find all proxies within a 1Mb window around each variant at R–squared thresholds of 0.1 and 0.8. We computed these from the UK10K^17^ sequence data on 3621 samples from two population cohorts (TwinsUK and ALSPAC) (data described elsewhere ^17^). Variants that were not observed in the UK10K data were excluded from our analysis.

### Data processing

Given a genome–wide distribution of p–values for association with a given disease or quantitative trait, we perform the following pre–processing steps in order to calculate the level of enrichment and its significance for an annotation of interest. To remove possible biases due to linkage disequilibrium (LD) or dependence between variants we compute the r^2^ between all SNPs within 1Mb windows and consider r^2^ of less than 0.1 between two variants to mean (approximate) independence. Next, from the full set of genetic variants for each phenotype, we create an independent set of SNPs where in order to keep all possible GWAS signals we sequentially find and retain the next most significant (lowest P–value) variant independent of all other variants in our independence set. After LD pruning an average of 6.1% (with range 5.5^-11^%) of genome–wide variants remained in our independence set for enrichment analysis (Supplementary Table S3). Next we annotate each independent SNP and consider it as overlapping a functional element if (i) the SNP itself resides in such a genomic region or (ii) at least one of its proxies in LD (r^2^>=0.8) and within 500Kb with it does. We include the latter as the association of a SNP in GWAS potentially tags the effect of other variants, which could underlie the observed association signal. The advantage of our greedy pruning over a P–value independent pruning is that we retain larger proportion of potentially causal variants (or tags of such SNPs). This is particularly advantageous for GWAS studies with low power and more pronounced at more stringent pruning thresholds.

### Quantifying enrichment

To find the enrichment of GWAS signals within a given functional annotation at a genome–wide significance P– value threshold T, we calculate the Fold Enrichment statistic as

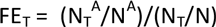

where N denotes number of all independent variants, N_T_– the subset of them with P–value less than T, N^A^– the number of variants in the annotation of interest, and N_T_^A^– the number of them with P–value less than T. This represents the ratio of proportion of variants in an annotation A at threshold T divided by the proportion of all variants at threshold T. We calculate FE at T=10^-1^, 10^-2^,…, 10^-8^ for all traits with at least 10 independent variants at a given threshold and test for significance at the four most significant GWAS cut–offs.

### Assessing statistical significance

To account for possible biases due to the GWAS P–value distribution depending on certain characteristics, which may also non–randomly associate with functional data, we use permutation testing, where we shuffle the P–values associated to each variant in our independence set in such a way as to match SNPs according to MAF, distance to nearest TSS and number of LD proxies (r^2^>=0.8) they have. Specifically, we bin variants according to 5 quantiles of MAF, number of LD proxies and distance to nearest TSS, resulting in 125 bins overall (default software options). We then permute variants within each bin separately. Due to the discreteness of the number of proxies and the skewness of their distribution in the pruned data, exact quantile binning is not always possible, in which case we create a stepwise binning in which we iteratively find the first (Q–q)’th quantile from the remaining variants after having already created q (out of Q) bins and removed those variants from consideration.

### Multiple testing

To account for multiple testing in the number of annotations used, we apply a Bonferroni correction for the number of independent tests carried out. Due to the nature of the data, annotations need not be (and are not in general) independent (e.g. biological replicates of the same cell types). Thus correcting for all annotations by assuming independence would be extremely stringent in practice. Instead, we estimate the effective number of independent tests performed similarly to Galwey, 2009 ^15^. More specifically, we take an independent subsample of SNPs and find the eigenvalues of the correlation matrix between all considered annotations and then find the effective number of independent test from equation 16 in Galwey, 2009. This results in at most 194 independent annotations out of a total of 424 for the DHS data (for the 27 phenotypes considered), to which we apply Bonferroni correction (p~2.5x10^-4^). Further details can be found in the supplementary material. Similarly for the segmentation data a total of 25x127=3175 annotations were used, which resulted in p~3.7x10^-5^ after correcting for multiple testing on the effective number of independent annotations at the 5% significance level.

### Number of permutations

For each annotation, we perform 10^5^ (DHSs) or 10^6^ (Segmentations) permutations to approximate the null distribution of our test statistic. These numbers are necessary in order to have sufficient resolution to detect significant enrichment results after multiple testing.

### Variability in enrichment significance

In order to assess the variability of our enrichment P–value estimates we performed enrichment analysis for Crohn’s Disease in DNaseI hypersensitive site data 1000 times for each of n=10^4^, 10^5^, 10^6^ and 10^7^ number of permutations per run. We then constructed 95% confidence intervals for–log10 enrichment P–value, where p=0 was denoted as equal to 1/n for these analyses.

### False positive rate

To get an estimate of GARFIELD’s false positive rate, we simulated ten phenotypes associated to variants selected at random from a pruned set of UK10K common variants (MAF>0.05). We selected between 5 and 100 variants in each case and used large effect sizes (1>beta>0.3) in order to ensure GWA analysis would detect those loci. We further simulated 1000 random annotations by mimicking the peaks lengths and between peak distances from the ENCODE HepG2 DHS cell line. We then performed enrichment analysis for each annotation–trait pair and estimated the false positive rate as the proportion of cell types showing significant enrichment for a given trait.

### Segmentation FE distribution and between trait sharing

From all significantly enriched cell types per trait and segmentation state, we calculated the median FE and then plotted its distribution across traits in order to estimate the per–state FE. Additionally, we counted the number of cell types per feature that were found to be significantly enriched and with FE>2 in a single trait, shared between traits with related function or shared between different classes of traits. For trait classification see Supplementary Table S3 or Table 1.

### Software

GARFIELD can be downloaded from http://www.ebi.ac.uk/birney–srv/GARFIELD/. The tool consists of two main parts: (i) pruning and annotation of the GWAS study of interest and (ii) calculating fold enrichment and testing its significance. Additionally, a quicker version of the method is implemented which terminates the draws for a given annotation if after a certain number of draws (minimum of 100 by default) it can no longer reach significance. In practice this is more advantageous for traits with few enrichments as opposed to ubiquitous ones. As well as this standalone software we have also developed an R package which can be downloaded from Bioconductor.

